# Expression of zebrafish *Brn1.2 (Pou3f2)* and two *Brn-3a* (*Pou4f1*) POU genes in brain and sensory structures

**DOI:** 10.1101/2021.05.26.445703

**Authors:** Satish Srinivas Kitambi, Gayathri Chandrasekar

## Abstract

POU genes are characterized by a conserved POU DNA-binding domain, and are divided into six subclasses. Class III and IV POU genes are predominantly expressed in the developing nervous system. POU class III genes are critical for several neuronal cell differentiation and class IV POU genes serve important functions in the differentiation and survival of sensory neurons. In this study, we attempted to identify POU genes in the zebrafish and pufferfish genomes by using existing bioinformatics tools. We analysed the expression of zebrafish *brn1*.*2* and *brn3a* genes (*brn3a1 and brn3a2)*) using whole-mount *in situ* hybridisation. Similarly to the mammalian orthologue, zebrafish *brn1*.*2* was widely expressed in the forebrain, midbrain and hindbrain. During the late stages of embryogenesis, *brn1*.*2* expressing cells were located in the preoptic area and in the auditory vesicles. Expression of both zebrafish *brn3a genes* was detected in trigeminal ganglia, cranial sensory ganglia, sensory neurons along the dorsal spinal cord, in the anterior and posterior lateral line placodes (ALL and PLL), retinal ganglion cell layer, optic tectum and in small cell clusters in the forebrain and hindbrain. Similar to mammalian Brn3a, zebrafish *brn3a* genes were detected in the retina and sensory structures. However, different domains of expression were also observed, namely in spinal sensory neurons, and lateral line system.

## Introduction

POU genes encode a subclass of sequence-specific DNA-binding proteins within the family of homeodomain transcription factors. They are defined by the conserved POU DNA-binding domain, that was originally identified by sequence comparison of the mammalian *Pit-1, Oct-1, Oct-2* homeodomain proteins and the *Caenorhabditis elegans* transcription factor *unc-86* (Herr et al., 1988). Based on their sequence homologies, POU genes have been divided into six subclasses (I to VI) (Latchman, 1999; Ryan and Rosenfeld, 1997). Mammalian class III POU genes include four members, *Brn-1, Brn-2, Brn-4* and *Oct-6/SCIP/Tst-1* that are predominantly expressed in the developing and adult nervous system (He et al., 1989; Monuki et al., 1989; Meijer et al., 1990; Hara et al., 1992; Mathis et al., 1992; Alvarez-Bolado et al., 1995; Zwart et al., 1996). POU class III genes are important for the neuronal cell differentiation. Knock out of the mouse *Brn -2* gene results in the loss of neurons that produce oxytocin, vasopressin and corticotropin-releasing hormone (Nakai et al., 1995; Schonemann et al., 1995). *Brn-4* mutants in mice show developmental defects in the inner ear causing deafness (Phippard et al., 1999). In humans, DFN3, an X-chromosome linked non-syndromic mixed deafness is caused by the naturally occurring mutations in *Brn-4* gene (de Kok et al., 1995). Targeted deletion of *Tst-1* has shown that *Tst-1* is essential for the terminal differentiation of myelinating Schwann cells in the peripheral nervous system (Bermingham et al., 1996; Jaegle et al., 1996).

In zebrafish, five POU class III genes have been identified and characterized (Matsuzaki et al., 1992; Sampath and Stuart 1996; Spaniol et al., 1996; Hauptmann and Gerster 1996; Hauptmann and Gerster 2000). They include *zp12, zp23, zp47, brn1*.*2* and *zp50*. The zebrafish POU class III genes show high sequence identity with their corresponding mammalian genes. Similar to their mammalian members, zebrafish class III POU genes lack introns within their POU-domain encoding sequences. z*p50* is orthologous to mammalian *Oct-6* (Levavasseur et al., 1998). *zp12* and *zp23* are the zebrafish orthologs of mammalian *Brn-1* and are identical with each other (Spaniol et al., 1996). *zp47* and *brn1*.*2* are related to each other and are the zebrafish orthologs of mammalian *Brn-2* gene (Spaniol et al., 1996; Sampath and Stuart 1996). The presence of additional genes in zebrafish could be attributed to the gene duplication in the genomic sequence of the teleost lineage. Previously only a partial sequence of *brn1*.*2* was deduced and a detailed developmental expression analysis of this gene was not described (Sampath and Stuart et al., 1996). Therefore, we have attempted to describe the developmental expression pattern of zebrafish *brn1*.*2* using whole-mount *in situ* hybridization.

Members of the class IV group of POU genes are characterized by an additional amino terminal consensus sequence, the POU-IV box (Gerrero et al., 1993; Xiang et al., 1995; Xiang et al., 1993) and are known to play important roles during development of the nervous system (Latchman, 1999; Ryan and Rosenfeld, 1997). Drosophila *I-POU* (*Acj6*) and *Caenorhabditis elegans unc-86* are the invertebrate homologues of mammalian class IV POU genes (Gruber et al., 1997). The three mammalian class IV POU genes, *POU4F1* (also called *Brn-3*.*0, Brn-3a, RDC1*), *POU4F2* (*Brn-3*.*2, Brn-3b*), and POU4F3 (*Brn-3*.*1, Brn-3c*) display high sequence similarities and distinct but overlapping expression patterns in the developing CNS and PNS (Latchman, 1999). Murine *Brn-3a* is mainly expressed in dorsal root and trigeminal ganglia, medial habenula, red nucleus and inferior olivary nucleus (Xiang et al., 1996). Loss of *Brn-3a* function by targeted deletion in mice causes loss of neurons in the brain stem and trigeminal ganglion and leads to uncoordinated limb movement and impaired suckling (Xiang et al., 1996). Human *Brn-3a* is localised on chromosome 13 and was found expressed in subsets of peripheral nervous system tumours (Collum et al., 1992). Human *Brn-3a* has been reported to activate expression of p53 in human tumour cells (Budhram-Mahadeo et al., 2002). It has also been shown that human Brn3a can activate the NGF1-A promoter in primary neurons and neuronal cell lines (Smith et al., 1999).

Zebrafish *brn3b* and *brn3c* homologues have recently been characterized, while a *brn3a* gene of zebrafish has not been described. In zebrafish two *brn3b* cDNAs, a long and a short isoform, have been cloned and their expression reported in the retina, optic tectum, migrating posterior lateral line primordium and larval neuromast (DeCarvalho et al., 2004). Zebrafish *brn-3c* has also been cloned and was found to be expressed in the developing otic vesicle (DeCarvalho et al., 2004; Sampath and Stuart, 1996). In an effort to determine the complete set of POU genes in teleost fish by search through available genome sequence and EST databases, we identified two *brn3a* zebrafish orthologues named *brn3a1* and *brn3a2*,. Similarly to *brn3b, brn3a2* was found to be expressed as a long (*brn3a2 (l)*) and a short (*brn3a2(s)*) isoform. The developmental expression patterns of the two-zebrafish brn3a genes were analyzed by whole-mount *in situ* hybridisation (WISH).

## 1. Results and discussion

### 1.1 Phylogenetic analysis of the zebrafish Pou gene class

Studies on vertebrate genome evolution provide increasing evidence of a whole genome duplication in the teleost lineage. In order to determine the effect of the proposed teleost genome duplication on the number of POU genes, we searched the zebrafish and pufferfish genome databases (Zebrafish version 3 and 4 (Zv3, Zv4) and FUGU 2.0) (www.ensemble.org). Our database search revealed 18 POU genes in zebrafish and 17 POU genes in pufferfish. To classify the identified POU genes into different subclasses, multiple sequence alignment was performed using the identified POU domain sequences and a phylogenetic tree was constructed (Fig 1A). The phylogenetic tree clustered the different zebrafish and pufferfish genes into the known six subclasses. Zebrafish and pufferfish genes corresponding to each of the six subclasses were identified. When compared to the mammalian set of 15 POU genes, it became eveident that several POU genes were present in two copies, while others were missing in the two teleost genomes. Surprisingly, genes corresponding to POU5F2 (Sprm-1) and POU3F4 (brn-4) could not be identified through the genome search in zebrafish and pufferfish. Perhaps, zebrafish and pufferfish sprm-1 and brn-4 may be secondarily lost during evolution (Fig 1A).

**Figure 1:**
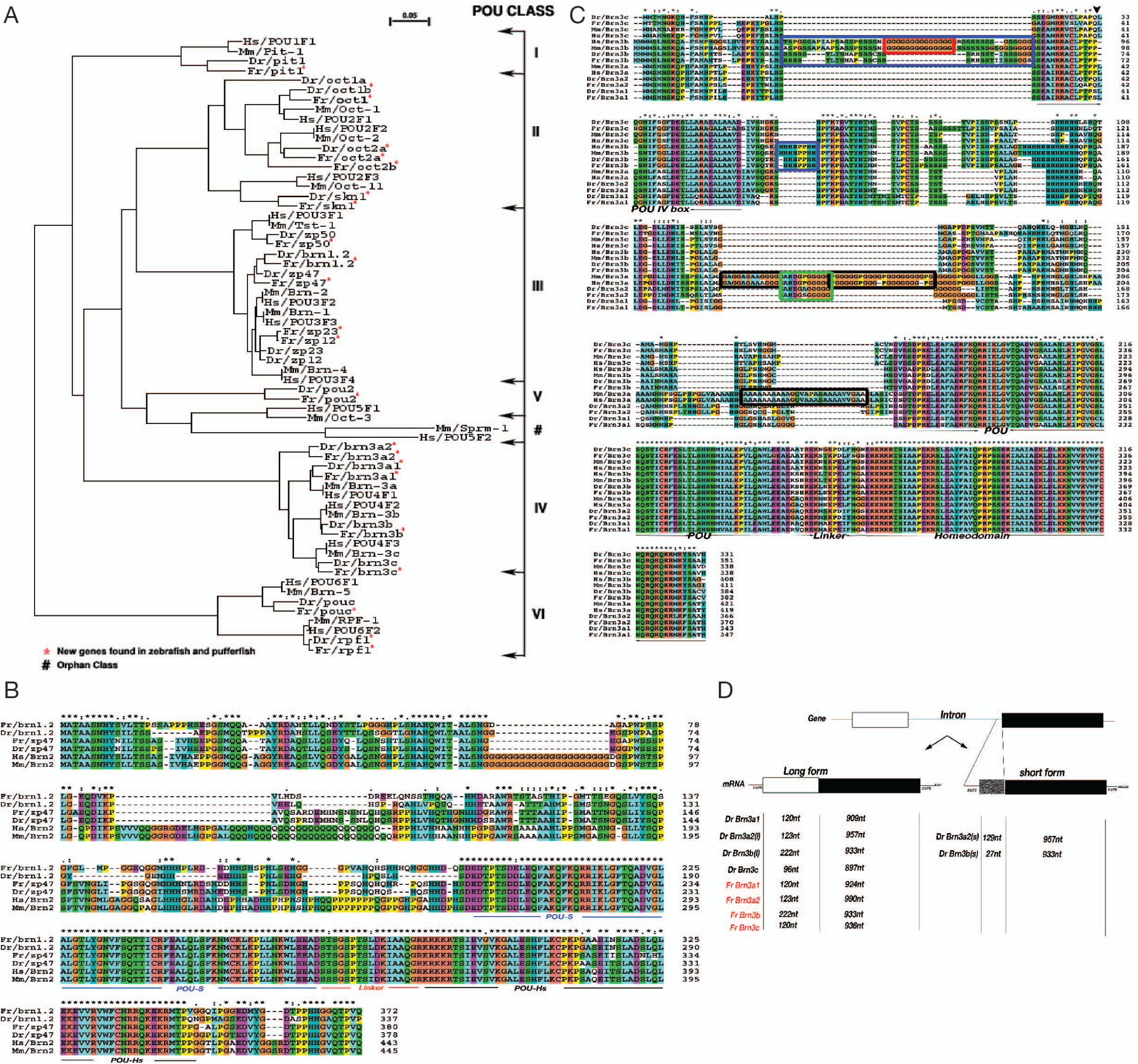
POU Family tree and sequence analyses of POU class III and IV. a) Phylogenetic tree showing the clustering of new zebrafish and pufferfish POU sequences into six different classes. Multiple sequence alignment of POU domain and phylogenetic tree was done with CLUSTAL X software and the phylogenetic tree was viewed with NJ plot software. Zebrafish *oct2b* could not be fully assembled and hence not included in the tree. B) Multiple sequence alignment of mammalian brn2 with zp47 and brn1.2 from zebrafish and fugu. The POU-specific (POU-S), linker and the POU-homeodomain (POU-Hs) are demarcated below the sequence and showed high degree of conservation. C) Multiple sequence alignment of POU class IV protein sequences. The arrowhead indicates the intron position. Arrows indicate POUIV box, POU specific, linker and homeobox domain region. The black box indicates region specific for mammalian brn3a class and the green box indicates region specific to mammalian Brn3a and fish brn3a2, brn3a1 of zebrafish and pufferfish lacks this region. The red box indicated the glycine rich region specific to mammalian Brn3b and the blue box indicates region specific for Brn3b. D) Zebrafish *brn3* family gene structure. The arrows indicate that all members of *brn3* form mRNA with two exons (lon form) and some of them form a shorter isoform. Zebrafish *brn3b* and *brn3a2* forms a longer (l) and shorter (s) isoforms, the full length coding sequence is made up of two exons and the shorter isoform is made up of one exon and a part of intron. The length of nucleotides (nt) of the exons and the part of the intron from different *brn3* sequences of zebrafish and fugu is indicated below.

### 1.2 Identification of zebrafish brn1.2 cDNA

We identified the zebrafish *brn1*.*2*, fugu *brn1*.*2* and *zp47* gene by TBLASTN search of the genomic zebrafish and fugu sequence at wwww.ensemble.org using zebrafish *zp47* (acc.no. P79746). The constructed sequence was used to screen the zebrafish EST database for zebrafish *brn1*.*2* ESTs. Two ESTS for zebrafish brn1.2 were identified and sequenced. We found that only one of them contained the complete open reading frame, therefore we used this clone for further analysis. This clone contained only a partial fragment of the 5ÚTR (20nt), 1639nt 3ÚTR and 1014nt ORF. Multiple sequence comparison between the zebrafish POU class III proteins Zp47 and Brn1.2 with the corresponding proteins of other species revealed high conservation in the POU domain region (Fig 1B). The POU domain sequence of *zp47* and *brn1*.*2* are highly similar to human, mouse and pufferfish. *zp47* showed an overall sequence identity of 97.4% to human, mouse and pufferfish while *brn1*.*2* was 95.5% identical to human and mouse and 96.8% identical to pufferfish.

### 1.3 Identification of zebrafish brn3a cDNAs

Through the zebrafish genome search ENSDARP00000007850 and ENSDARP00000013808 were identified. Similarity search with Blast program followed by multiple sequence alignment of POU domain and phylogenetic tree construction was performed to identify the class of POU genes these hits belonged. The similarity analyses revealed that ENSDARP00000007850 and ENSDARP00000013808 were paralogues to each other and orthologues to mammalian Brn3a.

For the murine *Brn3a* gene a short and a long cDNA isoform has been characterized (acc. Nos: AAO60105, AAO60106) (Thomas et al., 2004). In order to explore the possible existence of zebrafish *brn3a1* and *brn3a2* isoforms, we attempted to theoretically construct a long and a short isoform transcript form the available genome sequence using *in silico* methods. It was possible to construct a long and a short isoform for zebrafish *brn3a2 (brn3a2(l) and brn3a2(s))* but not for zebrafish *brn3a1*. The coding region of zebrafish *brn3a2(l)* was derived from two exons (E1,E2) separated by an intron (I1). Incontrast, *brn3a2(s)* was made up from the 3’ region of intron (I1) and exon E2 (Fig 1D). The genome sequence analyses of the members of the *brn3* family in zebrafish and fugu demonstrated that the mRNA was mainly derived from two exons and some members of the family also formed shorter isoforms (Fig 1D).

The amino acid sequences from the assembled zebrafish *brn3a1, brn3a2(l)* and *brn3a2(s)* were taken to mine the zebrafish expressed sequence tag database available at NCBI (www.ncbi.nlm.nih.gov). One EST for zebrafish *brn3a1* was identified and sequenced. The sequence contained 165 nt 5’ UTR and 1133 nt 3’UTR and an open reading frame encoding 344 amino acids. Two EST’s for zebrafish *brn3a2(l)* were identified and sequenced. The sequence data revealed an open reading frame of 366 amino acids and 17nt 5’UTR and 1254nt 3’UTR. Similarly one EST for *brn3a2(s)* was identified and sequenced revealing an open reading frame of 367 amino acid and 7nt 5’UTR and 1254nt 3’UTR. The sequences of all Ests matched the *in silico* sequence prediction from the zebrafish genome. Similar *in silico* sequence analyses of pufferfish genome lead to the identification and construction of pufferfish *brn3a1, brn3a2(l)* and *brn3a2(s)* sequences. Sequence Identity of the POU domain region between the zebrafish brn3a genes and mammalian Brn3a was very high (Human and Mouse Brn3a is 93.7% identical to zebrafish brn3a1 and 96.2% identical to zebrafish brn3a2). Even higher sequence identity was found between zebrafish and pufferfish class IV POU domain sequences (pufferfish brn3a1 is 97.5% identical to zebrafish brn3a1 and pufferfish brn3a2 is 97.5% identical to zebrafish brn3a2).

### 1.4 Multiple sequence alignment of Pou class IV sequences

Multiple sequence alignment of class IV amino acid sequences revealed a high conservation of the POU domain region (POU+Linker+Homeodomain) between different species, but significant conservation was also observed outside of the POU domain (Fig 1C). A poly-glycine stretch at the N-terminus of mammalian Brn-3b and in mammalian Brn-3a was seen; this region was absent in the zebrafish and pufferfish sequences. Distinct region specific for brn3b sequences, mammalian brn3a, brn3b sequences were observed (Fig 1C). A specific region of 11 amino acid was identified to be present in fish (zebrafish and pufferfish) brn3a2 and mammalian brn3a. This region was absent from the fish brn3a1 (Fig 1C). Analysis of the genomic sequence of *Pou class IV* genes from human, mouse, zebrafish, and pufferfish showed that the coding region is derived from two exons. The position of the intron was conserved among all the species compared. The length of the exons of different brn3 genes in zebrafish and fugu were very similar and zebrafish brn3a2 and brn3b also formed shorter isoforms (Fig 1D).

### 1.5 Developmental expression pattern of zebrafish brn1.2

We performed whole-mount *in situ* hybridization (WISH) (Hauptmann and Gerster, 1994, 2000) to characterize the spatial expression of *brn1*.*2* in the developing zebrafish brain. Expression of *brn1*.*2* was first detected at the tail bud stage in an area that corresponds to the midbrain primordium (Fig 2A). At the 3-, 5- and 10-somite stages the expression became quite strong in the midbrain region. From the 5-somite stage on *brn1*.*2* was also detected in the hindbrain and spinal cord (Fig 2B,C,D).

**Figure 2:**
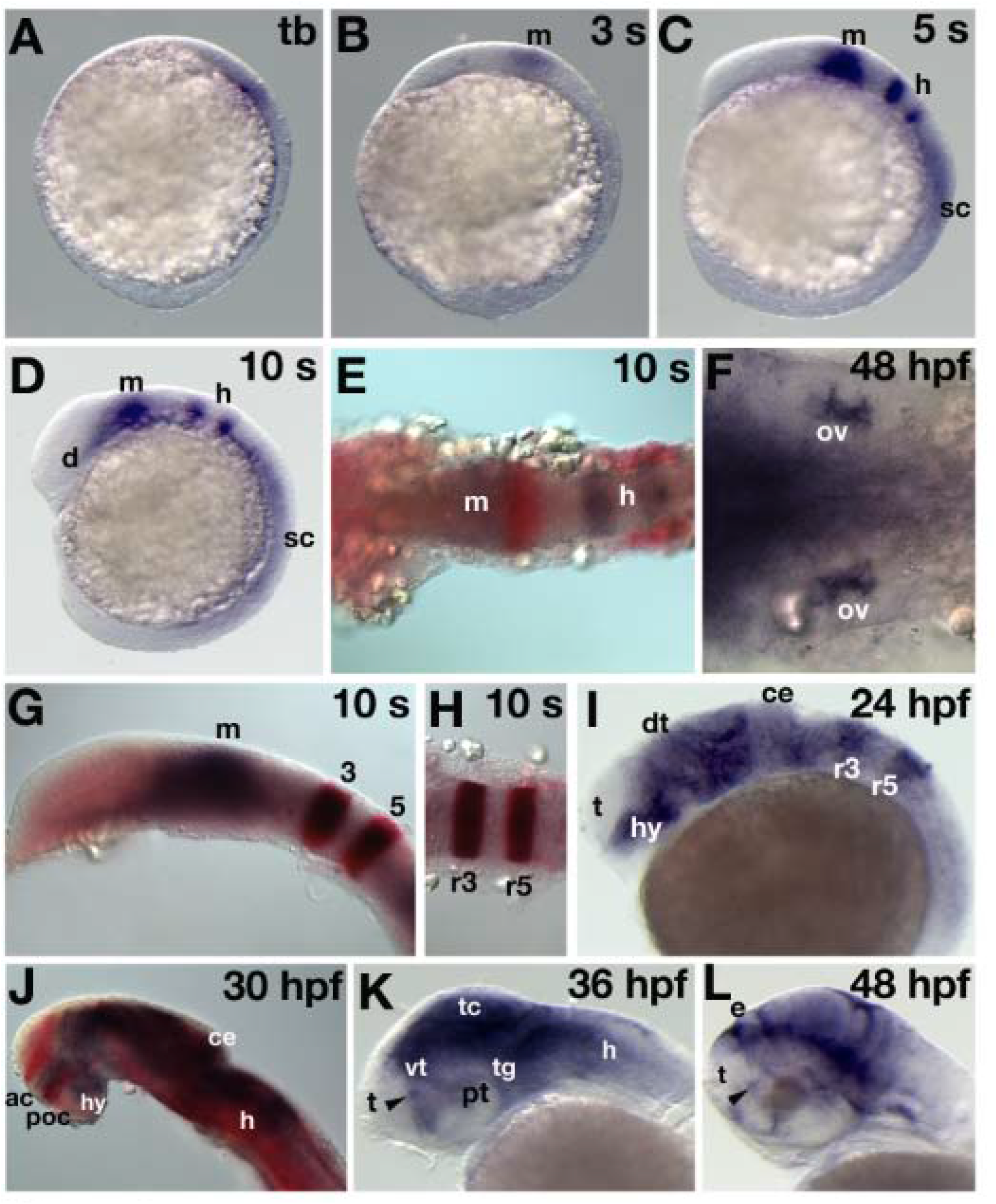
Expression of *brn1*.*2* in the developing zebrafish brain. Embryos were subjected to WISH to visualize *brn 1*.*2* in blue, *pax2*.*1* in red (E) and *krx-20* in red (G, H). Embryo in panel J was subjected to immunohistochemistry to visualize the primary axons in red after detecting *brn1*.*2* in blue via WISH. Panels A, B, C, D, G, I, J, K, L show lateral views with the anterior to the left. Panels E, F, H show dorsal views with the anterior to the left. Developmental stages are indicated on the right hand corner of each panel. Arrowhead in panel K and L indicate the preoptic region of the brain. Abbreviations: ce, cerebellum, dt, dorsal thalamus, h, hindbrain, hy, hypothalamus, m, midbrain, ov, otic vesicle, pr, pretectum, r3, rhombomere 3, r5, rhombomere 5, sc, spinal cord, tc, tectum, tg, tegmentum, t, telencephalon, vt, ventral thalamus.

A weak expression of *brn1*.*2* transcripts was found in the ventral diencephalic region at the 10-somite stage (Fig 2D). We compared the expression of *brn1*.*2* with *pax2*.*1* (Krauss et al., 1991) and *krx-20* (Oxtoby and Jowett, 1993) in order to locate the position of *brn1*.*2* transcripts in the midbrain and hindbrain. Two color *in situ* hybridisation with *pax2*.*1* showed that *brn1*.*2* was located anterior to *pax2*.*1* expression domain indicating that *brn1*.*2* expression domain was located in the anterior midbrain (Fig 2E). Co-labelling studies with *krx-20* revealed that *brn1*.*2* expression in the hindbrain is confined to r3 and r5 (Fig 2G,H).

From 24hpf on, *brn1*.*2* showed a widespread expression in the CNS. In the forebrain, *brn1*.*2* was detected in the diencephalon, epiphysis, ventral thalamus, dorsal thalamus and pretectum while the telencephalon was devoid of *brn1*.*2* expression. At 1dpf, *brn1*.*2* was expressed throughout the hindbrain with strong expression levels in r3 and r5 (Fig 2I). At 30hpf a lamda shaped expression domain with two arms were observed in the diencephalon (Fig 2J). To define the position of the two arms more precisely we performed whole-mount *in situ* hybridisation to visualize *brn1*.*2* followed by immunohistochemistry to detect acetylated α-tubulin (Piperno and Fueller, 1985). One arm of the lamba domain was found to be lying in-between the optic recess and the tract of the postoptic commissure. The other arm extending from the ventral thalamus was broader in shape and was widespread across the tract of the commissure of the posterior tuberculum (Fig 2J).

At 36hpf and 2dpf the expression of *brn1*.*2* became more complex in the brain (Fig. 2K,L). At 2dpf, few *brn1*.*2* positive cells were found scattered in the telecephalon (Fig. 2L). Additionally, *brn1*.*2* expression was detected for the first time in the auditory vesicles at this stage of development (Fig. 2F). Interestingly, at 36hpf and 48dpf few cells expressing *brn1*.*2* were concentrated in the preoptic area of the forebrain (Fig. 2K & L). The mammalian neuro-hypophyseal hormones oxytocin and arginine vasopressin are represented as isotocin and vasotocin in fish. Two color *in situ* hybridization with zebrafish isotocin-neurophysin (*itnp*) and vasotocin-neurophysin (*vt*) was performed to determine whether *brn1*.*2* was co-expressed with *itnp* and *vt*. In zebrafish, *itnp* mRNA is expressed as bilateral cell clusters in the dorsal preoptic region of the brain (Unger and Glasgow, 2003). Zebrafish *vt* is first detected at 24hpf in the anterior diencephalon (not shown). At 48 hpf, *vt* is expressed as bilateral cell clusters in the preoptic area and in the ventral hypothalamus (Fig. 3E). Co-labelling experiments performed on 48hpf embryos revealed that *brn1*.*2, itnp* and *vt* expressing cells are located in the same region in the preoptic area (Fig. 3A-F). In a separate study it was demonstrated that the isotocin expressing cells are co-distributed with corticotropin-releasing hormone (CRH) producing cells in the preoptic region of the forebrain (Fig. 3G-I) (Chandrasekar et al., 2007). Therefore it is likely that the CRH expression domain lies within the *brn1*.*2* expression domain in the preoptic area. In mammals, so far, four transcription factors, namely, *Sim1, Arnt 2, Otp* and *Brn-2* have been known to be required for the development of specific neuroendocrine cell types located within the paraventricular nuclei (PVN) and supraoptic nuclei (SON) of the hypothalamus. *Sim1* and its heterodimerization partner *Arnt2* belong to the bHLH-PAS family of transcription factors. Both *Sim1* and *Arnt2* are critical for the development of the neurosecretory cell types that produce oxytocin (OT), arginine vasopressin (AVP), corticotropin-releasing hormone (CRH), thyrotropin-releasing hormone (TRH) and somatostatin (Keith et al., 2001; Michaud et al., 2000; Michaud et al., 1998). *Brn-2* is the downstream target of *Sim1* and is required for the terminal differentiation of OT, AVP and CRH expressing neurons (Nakai et al., 1995; Schonemann et al., 1995). *Brn-2* also regulates the transcription of CRH by binding to its promoter (Li et al., 1993). The homeobox gene *orthopedia* (*Otp*) function in parallel with *Sim1* and both are required for the maintenance of *Brn-2* expression (Acampora et al., 1999). A recent study in zebrafish shows that *sim1* and *otp* are required for isotocin cell development and they act in parallel to direct the differentiation of isotocin cells in zebrafish (Eaton and Glasgow, 2006; Eaton and Glasgow, 2007). Considering our results from the co-expression studies we are tempted to speculate that similar to the mammalian *Brn-2* gene, zebrafish *brn1*.*2* could also have a role in the differentiation of isotocin, vasotocin and CRH producing cell types.

**Figure 3:**
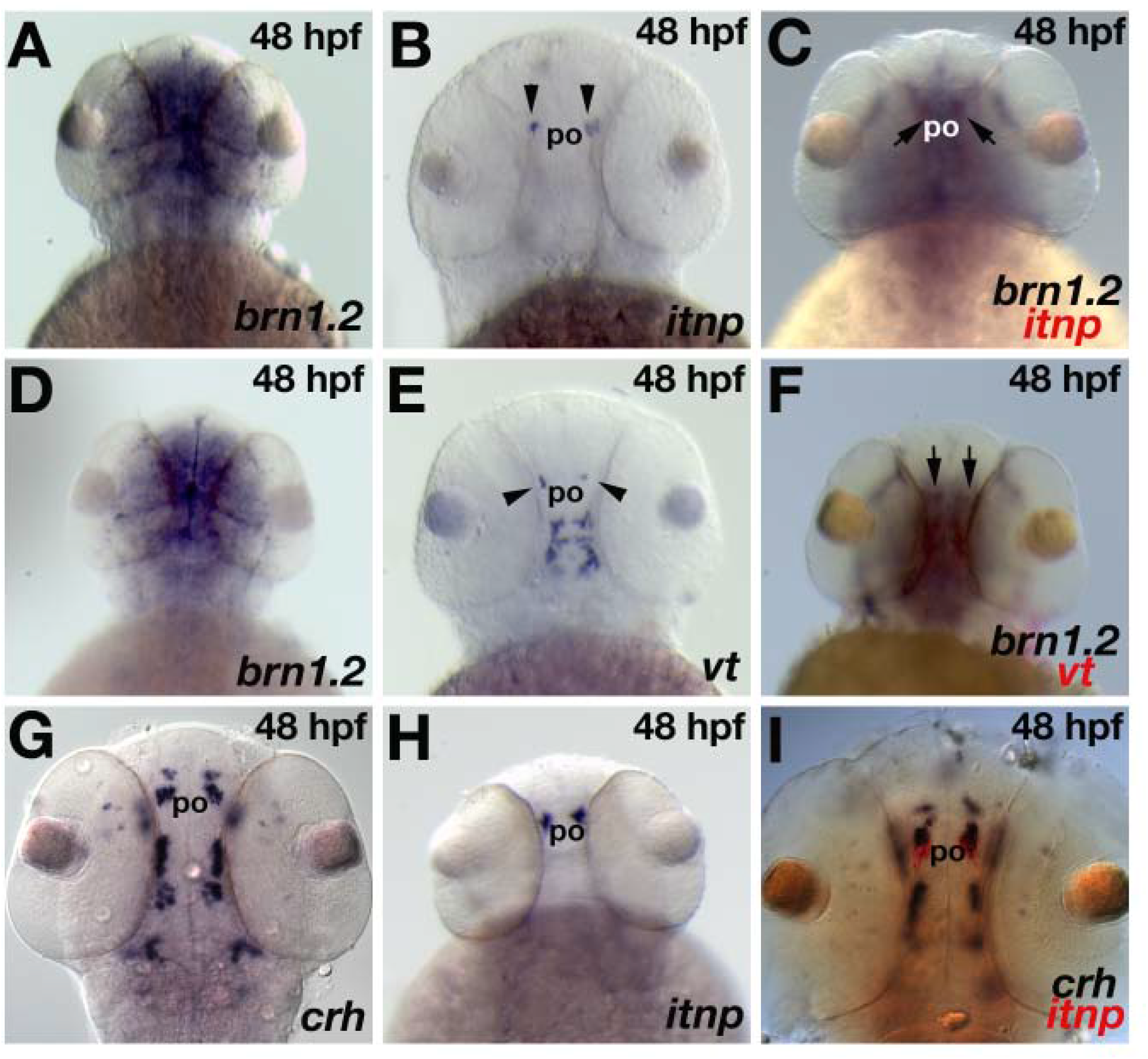
Localization of *brn1*.*2* in the preoptic area of the hypothalamus. Two-color *in situ* hybridization was performed to examine co-expression of *brn 1*.*2* with *itnp* and *vt* and also *crh* with *itnp*. Embryos were subjected to WISH to visualize *brn 1*.*2* in blue (A, C, D, F), *itnp* in blue (B, H) and in red (C, I), *vt* in blue (E) and in red (F) and *crh* in blue (G, I). Dorsal views of 48hpf embryos are shown (A-I). Anterior is facing up. *itnp* and *vt* expression domain in the preoptic area are indicated by arrowheads (B, D). Arrows in (C, F) indicates the localization of *itnp* and *vtnp* within *brn1*.*2* expression domain in the preoptic area. Abbreviations: po, preoptic area.

In conclusion, we have analyzed the developmental expression pattern of *brn1*.*2* in the embryonic brain. Our results show that zebrafish *brn1*.*2* exhibits a very complex expression pattern in the developing brain very much similar to its paralog *zp47* (Hauptmann and Gerster, 2000a). Overlapping expression domains of *brn1*.*2, itnp* and *vtnp* in the preoptic area suggests a possible role for *brn1*.*2* in the development of specific neuroendocrine cell types.

### 1.6 Expression of zebrafish brn3a genes during development

Whole-mount *in situ* hybridization was used to study the developmental expression profile of *brn3a1* and *brn3a2*. Fluorescein labelled *neuroD* was used to identify the sensory structures (Blader et al., 1997; Kim et al., 1997; Korzh et al., 1998). Initial expression of *brn3a1* was detected at the tailbud stage. *brn3a1* expression was confined to the trigeminal placode and to longitudinal stripes along the lateral aspects of the neural plate, presumably Rohon beard sensory neuron precursors (Fig. 4A-C, 5H). This expression pattern stayed essentially the same up to the 5 somite stage (Fig. 4D-I). From the 7 somite stage onwards to 15 somite stage, *brn3a1* expression was also found in the forming anterior and posterior lateral line placodes (ALL and PLL) (Fig. 4J,K,M,N, 5I) in addition to the longitudinal stripes along the developing spinal cord. At 25 hpf, *brn3a1* expression had spread to various cranial sensory placodes, including anterior dorsal (AD), anterior ventral (AV), facial (F), middle lateral line placode (M) and posterior lateral line placode (PLL) Fig. (5A-C) and remained unchanged at 30 hpf and 36h pf (Fig 5F, G, J, K). By 48hpf, *brn3a1* expression in cranial ganglia was decreased and restricted to the neuromast of ALL and PLL (Fig 5L). Apart from these regions, *brn3a1* was also expressed in cluster of cells in the forebrain and hindbrain (fig. 5J). The expression of *brn3a1* in Rohan Beard neurons was transient and disappeared by 30 hpf.

**Figure 4:**
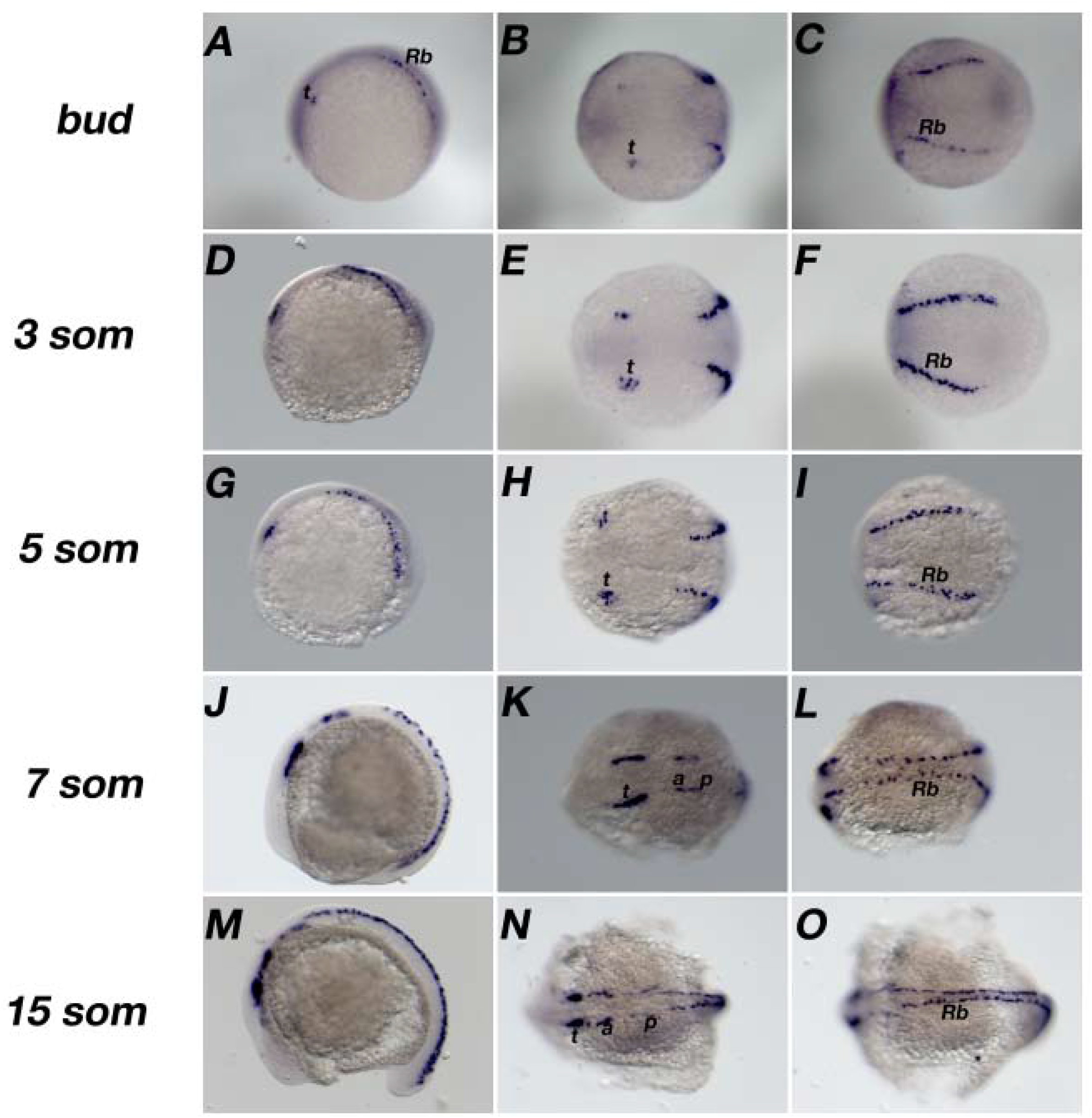
Early expression of zebrafish *brn3a1*. Anterior to the left. Lateral (A,D,G,J,M) and dorsal (B,C,E,F,H,I,K,L,N,O) views are shown. Expression is observed in Rohon beard neurons (Rb) and in the trigeminal (t), anterior lateral line ganglia (a), posterior lateral line ganglia (p).

**Figure 5:**
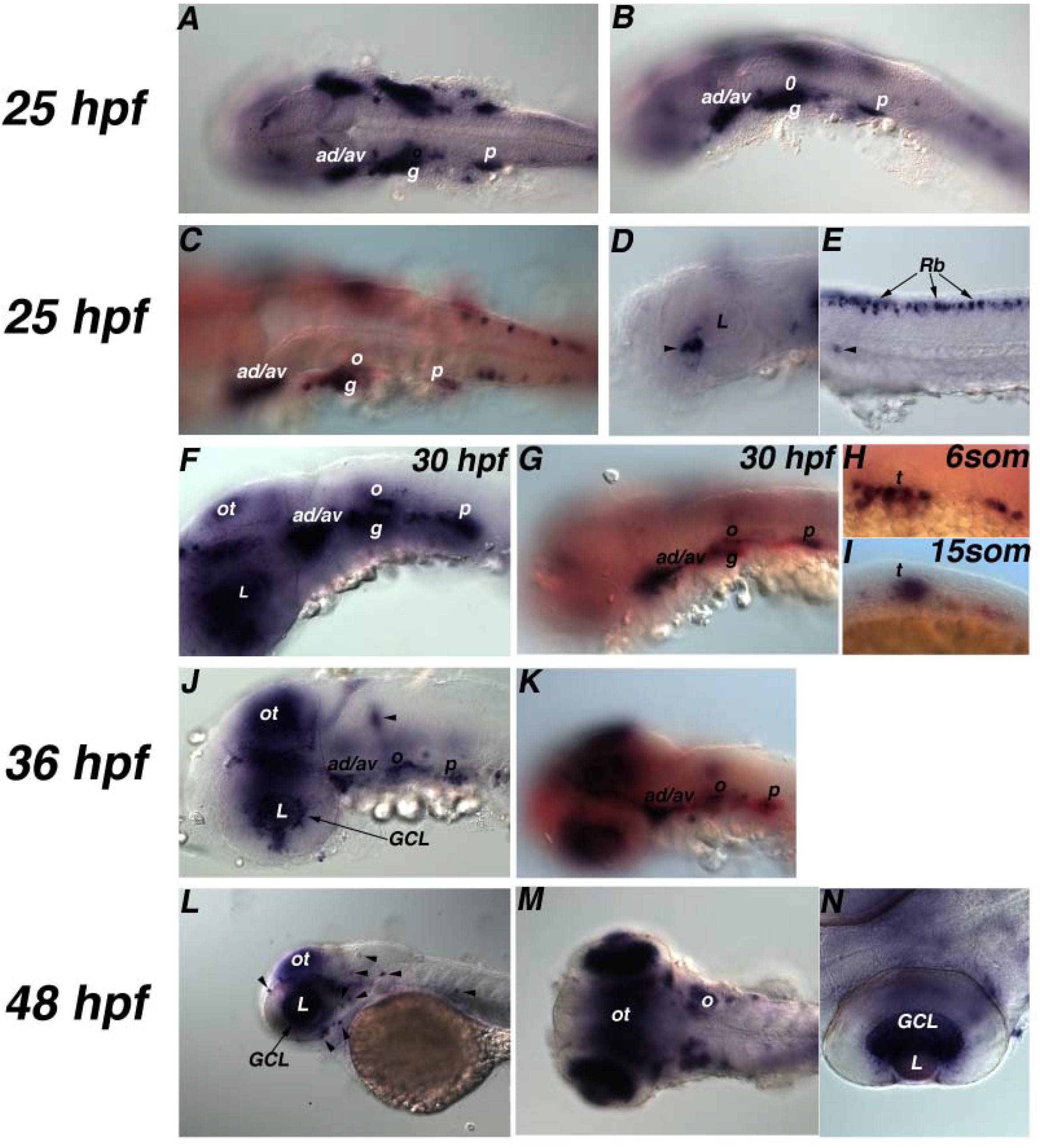
Late spatial expression of zebrafish *brn3a1*. In all panels, anterior is to the left. Lateral (D, E, F, H, I, J, K, L), dorsal (A, M, N) and oblique lateral (B, C, G) views of embryos are shown. Expression is seen in the anterior dorsal (ad), anterior ventral (av), glossopharyngeal (g), posterior lateral line (p), rohan beard neurons (Rb), optic tectum (ot), ganglion cell layer (GCL),. Images H and I show expression in trigeminal ganglia (t), Arrowheads indicate expression in ventronasal region (image D), cluster of hindbrain cells (image J), neuromast (image L). Zebrafish *brn3a1* is stained in blue and zebrafish *neuroD* in red (C, G, H, I, K).

The retinal ganglion cells (RGC) are the first neurons to become post mitotic (Brown et al., 2001; Hu and Easter, 1999; Kay et al., 2001; Wang et al., 2001). The laminar pattern of the eye is formed by 60hpf (Hu and Easter, 1999) and the RGC’s are the first neurons to be born (Brown et al., 2001; Hu and Easter, 1999; Kay et al., 2001; Wang et al., 2001) and require *Ath5* expression for its determination (Brown et al., 2001; Kay et al., 2001; Liu et al., 2000; Liu et al., 2001; Wang et al., 2001). In mammals *Brn3a* is expressed in the RGC cells (Xiang et al., 1996). Zebrafish *brn3a1* was found to be expressed in the ventronasal region of the retina at 25 hpf (Fig 5D). Retinal *brn3a1* expression spread dorsally and temporally and throughout the RGC layer by 36hpf (Fig. 5J). The three retinal layers (ganglion cell layer, inner nuclear layer and outer nuclear layer) are clearly visible at 48hpf, and intense expression of *brn3a1* was seen throughout the RGC layer (Fig 5M-N). The early expression profile of zebrafish *brn3a1* indicated that *brn3a1* may precede the expression of *brn3b* and *brn3c*. The sequence of expression of *Brn3a, Brn3b* and *Brn3c* is different in mouse where the expression of *Brn3b* precedes the expression of *Brn3a* and *Brn3c* (Xiang, 1998). Axonal projections from the RGC’s exit the retina by 34hpf and innervate optic tectum by 72hpf (Burrill and Easter, 1995). Expression of *brn3a1* was also found in the optic tectum. Tectal expression of *brn3a1* was detected at 30 hpf (Fig 5F) and fully established by 48 hpf (Fig 5L-M), Zebrafish brn3a2 was very similarly expressed as brn3a1.

Expression of *brn3a2(l)* in the trigeminal placode and presumed Rohon Beard sensory neurons started around the 7 somite stage (Fig. 6A,B) and became stronger by 10 somite stage (Fig 6C,D). Expression in the trigeminal ganglion ceased at 24 hpf. Expression of *brn3a2(l)* was detected in the anterior lateral line (ALL) and posterior lateral line (PLL) by 15 somites and continued to be expressed there until 48hpf (Fig 6E-J, 7A-M). Clusters of cells expressing *brn3a2(l)* were also seen in the forebrain, hindbrain (Fig 7A,D,G,I) and along the dorsal cells of the spinal cord (Fig 7C,F). The expression in the spinal cord goes down by 48hpf (Fig 7M). In the midrain tectum and RGC layer, brn3a2*(l)* expression was established by 48hpf (Fig 7I,J,L,M). In summary, apart from expression of both *brn3a* genes in the RGC layer and sensory ganglia, expression of these genes was also detected in sensory neurons along the dorsal spinal cord and in small cell clusters within the forebrain and hindbrain. Similar to zebrafish *brn3b* (DeCarvalho et al., 2004), *brn3a1* and *brn3a2(l)* were also detected in the lateral line system.

**Figure 6:**
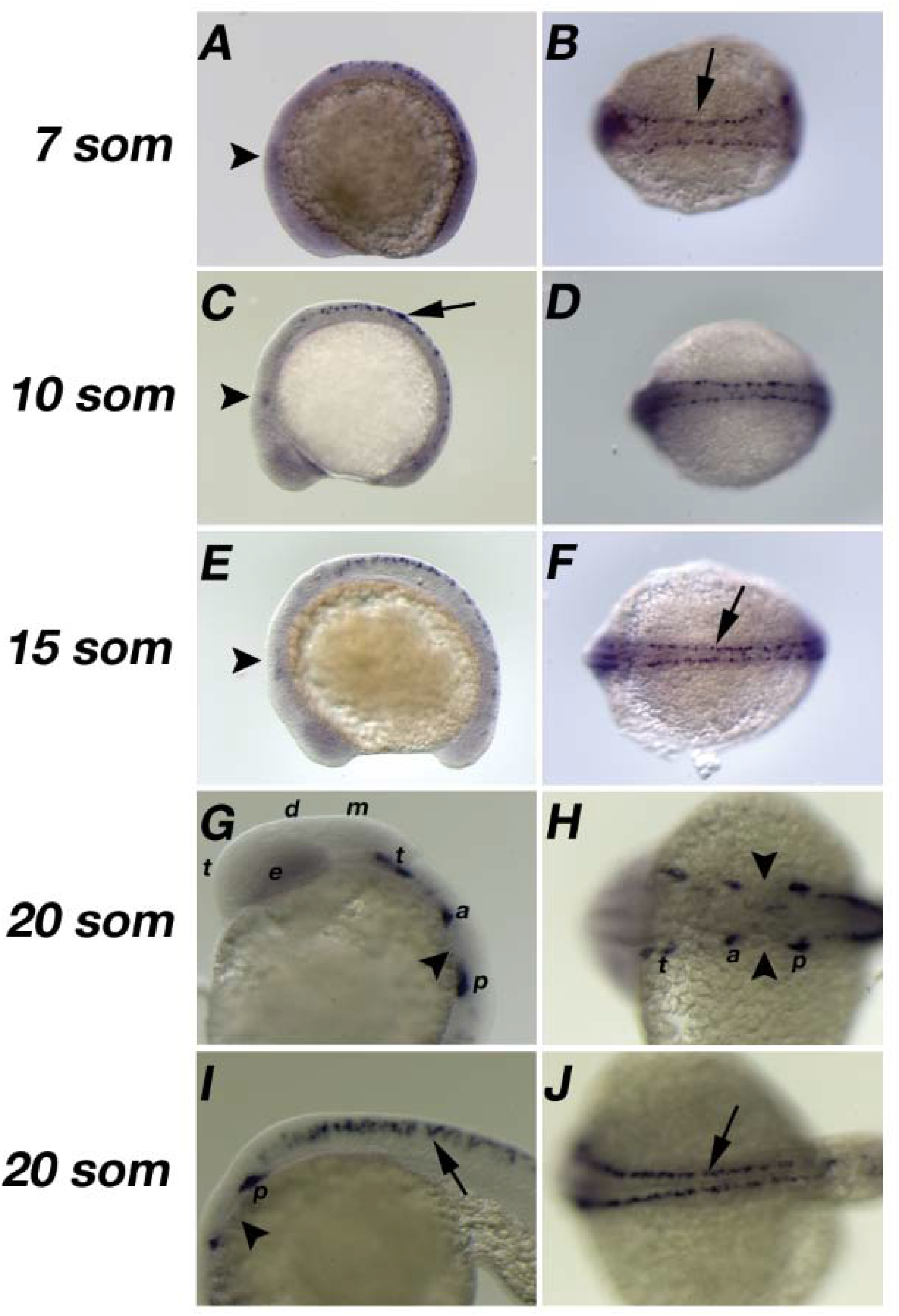
Expression of zebrafish *brn3a2(l)* from 7 to 20 somite stage. Images present lateral (A, C, E, G, I) and dorsal (B, D, F, H, J) views of the embryos. The figures are arranged with anterior to the left. Arrowheads indicate trigeminal ganglia (A, C, E) and otic placode (G, H, I). Arrows mark dorsal cells of the spinalcord. Expression is seen in the trigeminal ganglia (t), dorsal cells of the spinal cord (represented by arrows), anterior lateral line (a), posterior lateral line (p). Eye is represented by (e).

**Figure 7:**
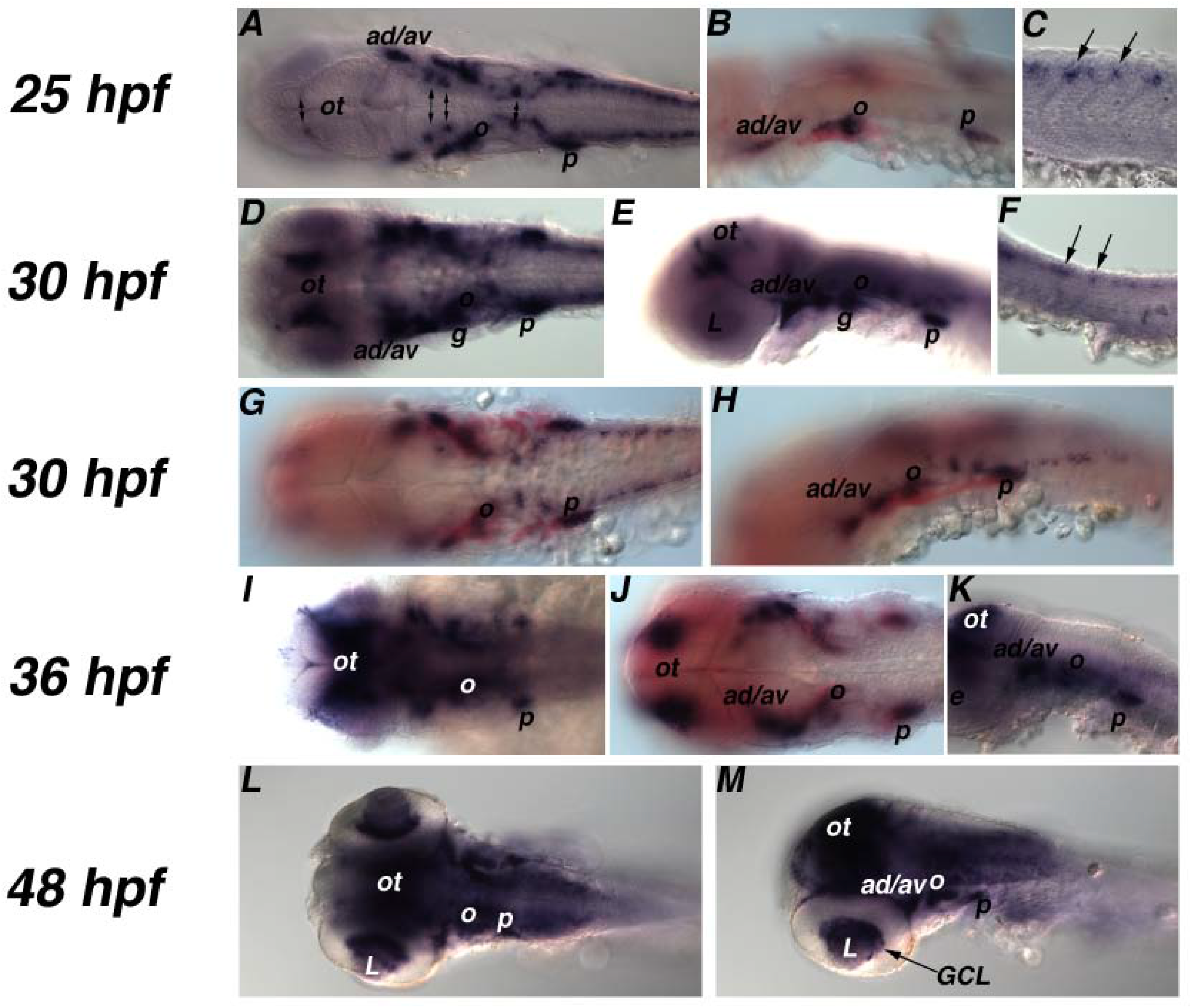
Expression of zebrafish *brn3a2(l)* from 25 hpf - 48 hpf. Embryos are arranged with anterior to the left. Expression is seen in anterior dorsal (ad), anterior ventral (av), otic (o), glassopharyngeal (g), posterior lateral line (p), optic tectum (ot), ganglion cell layer (GCL), dorsal cells of the spinal cord (represented by arrows), cluster of cells of the fore-, mid-, and hind brain (represented by double sided arrows). Lense of the eye is represented by (L). Expression of zebrafish *brn3a2(l)* is visualized in blue and zebrafish *neuroD* is shown in red.

## 2. Experimental procedures

### 2.1 Zebrafish maintenance

Wild type Zebrafish were maintained at 28.5 °C and under standard conditions of feeding, care and egg collection. Embryos were collected by natural mating. The collected embryos were staged according to Kimmel et al. (1995). Embryos were staged in hours post fertilization (hpf) and days post fertilization (dpf), and embryo stages older than 24hpf were subjected to 0.03% phenylthiourea treatment. The collected embryos were fixed at different stages in 4% paraformaldehyde overnight and then washed with phosphate buffered saline containing 0.1% Tween-20 (PBSTw) and stored in 100% methanol until usage for *in situ* hybridization.

### 2.2 Sequence analysis

Zebrafish and Pufferfish genome sequence databases were mined to identify and construct POU class genes. Mammalian POU genes were used as query. A database using Filemaker Pro software was developed to collect and maintain the assembled POU sequences from zebrafish and pufferfish as well as published sequences from other species. The assembled sequences were translated and POU, Linker region and Homeodomain were extracted and multiple sequence alignments were performed with the Clustal X program (Thompson et al., 1997). A neighbour joining tree with bootstrap value of 1000 was constructed based on the multiple sequence alignment obtained in the Clustal X program. The neighbour joining tree helped to cluster the newly found zebrafish and pufferfish sequences to their mammalian counterparts. The tree was viewed with NJ plot software (Perriere and Gouy, 1996).

Zebrafish *brn1*.*2, brn3a1* and *brn3a2* genomic sequences (ENSDARG00000023662, ENSDARP00000007850 and ENSDARP00000013808) were identified using zebrafish zp47 for brn1.2 and murine Brn3a protein sequences (acc No P79746 and acc. Nos. S69350) as query against the ongoing zebrafish genome project (Zv3) using TBLASTN search. The long and the short isoforms of *brn3a2* were assembled manually using conceptual translation. Amino acid sequences of *brn3a1, brn3a2(l), brn3a2(s)* were obtained by conceptual translation. The constructed zebrafish protein sequence of *brn1*.*2, brn3a1, brn3a2(l)* and *brn3a2(s)* were used as query to identify corresponding cDNAs from the zebrafish EST database. The database search revealed two EST’s for brn1.2 (LLKMp964N034Q2, IMAGp998L14324Q3), two EST’s for *brn3a1* (IMAGp998K1214311Q3, IMAGp998I1314318Q3), one EST for *brn3a2(l)* (IMAGp998P1314603Q3) and one EST for *brn3a2(s)* (IMAGp998A1414316Q1). The ESTs were purchased from the German Resource Center for Genome Research (www.rzpd.de), fully sequenced by MWG and submitted to GenBank (acc. Nos.).

The zebrafish Brn3a1 and Brn3a2 protein sequences were compared to the Brn3 sequences of other species. Multiple sequence alignments were performed with the Clustal X program (Thompson et al., 1997). The accession numbers for the used sequences were as follows: AAU13951, NP_620395, NP_620394 of (Mm) Mus musculus, NP_006228, NP_002691, NP_004566 (Hs) Homo sapiens, NP_571353, NP_997972, ENSDARP00000007850, ENSDARP00000013808 (Dr) Danio rerio, SINFRUP00000143813, SINFRUP00000162286, SINFRUP00000133796, SINFRUP00000133155 (Fr) Fugu rubripes.

### 2.3 Whole-mount in situ hybridisation (WISH)

Single color and two-color WISH was performed as described previously (Hauptmann and Gerster, 1994; 2000). *brn1*.*2, brn3a1* and *brn3a2(l)* specific digoxigenin-labeled antisense riboprobes were synthesized by KpnI (ASP718) linearization and T7 polymerase transcription. *brn1*.*2, brn3a1* and *brn3a2(l)* transcripts were visualized with anti digoxigenin-alkaline phosphatase conjugates and BCIP/NBT or BM purple substrates (Roche). For two color WISH *krx-20* (Oxytoby and Jowett, 1993), *neuroD* (Blader et al., 1997; Kim et al., 1997; Korzh et al., 1998) and *pax2*.*1* (Krauss et al., 1991) fluorescein labelled probe was used. The fluorescein labelled probe was detected with anti-fluorescein-alkaline phosphatase conjugates and Fast Red substrate. A Zeiss Axioplan DIC compound microscope and a Leica MZ16 dissecting microscope were used to view the embryos. Embryos were photographed with a Zeiss Axiocam digital camera and Leica DFC300FX digital camera. Image processing and composite figures were assembled with Adobe Photoshop 7.0 software.

### 2.4 Immunohistochemistry

Some embryos processed for WISH to visualize *brn1*.*2* were further processed for immunohistochemical detection of the position of the primary axons using a monoclonal antibody against acetylated alpha-tubulin (Piperno and Fuller, 1985). The experiments were performed as described previously (Hauptmann and Gerster, 1996). Axon tracts were visualized in red using Fast red as alkaline phosphatase substrate.

## Acknowledgements

We thank School of Lifesciences, Sodertorns Hogskola, Department of Biosciences, Karolinska Institutet, Institute for Healthcare Education and Translational Science (IHETS) and Kitambi Foundation for financial support. Dr. Giselbert Hauptmann for providing useful inputs into this study.

